# Foraging Ants as Liquid Brains: Movement Heterogeneity Shapes Collective Efficiency

**DOI:** 10.1101/2025.02.28.640754

**Authors:** Pol Fernández-López, Daniel Oro, Roger Lloret-Cabot, Meritxell Genovart, Joan Garriga, Frederic Bartumeus

**Affiliations:** Theoretical and Computational Ecology Group, Centre d’Estudis Avançats de Blanes (CEAB-CSIC), Cala Sant Francesc, 14, Blanes 17300, Spain; Centre de Recherches sur la Cognition Animale (UMR5169), Centre de Biologie Intégrative, Université de Toulouse, CNRS, UPS, Toulouse 31062, France; Institució Catalana de Recerca i Estudis Avançats, ICREA, Passeig Lluís Companys, 23, Barcelona 08010, Spain; Centre de Recerca Ecològica i Aplicacions Forestals (CREAF), Cerdanyola del Vallès, Barcelona 08193, Spain

## Abstract

Liquid brains conceptualize living systems operating without central control, where collective outcomes emerge from local but dynamic interactions. Therefore, movement is expected to shape the connectivity among individuals, allowing the system to optimize its efficiency. We empirically measured ant movement behavior across large spatiotemporal scales, closely reflecting the ecology of our model species, *Aphaenogaster senilis*. We then incorporated this into a liquid brain framework, enabling a quantitative replication of ant foraging efficiency and their spatiotemporal dynamics. Our results highlight that a simple feedback mechanism explains the foraging patterns of this species. Indeed, such feedback is modulated by adjusting the proportion of two coexisting movement behaviors: while the recruits facilitated information transfer and food exploitation by aggregating closely to the nest, the scouts mostly bypassed this feedback, enabling the discovery of alternative food sources. These findings underscore how complex systems frameworks can benefit from empirical insights, enhancing our understanding of the mechanisms underlying collective intelligence in biological systems.

## Introduction

Complex systems frameworks are relevant for understanding how natural systems operate without central control, manage numerosity, and show adaptive behavior [1, 2]. These frameworks offer valuable insights into the fundamental principles of cognition and organization in nature, such as what is observed in social species. However, said approaches often depend on simplifications and metaphors that may not fully represent the reality of living systems [3]. This is especially true in ecological systems, whose dynamics are conditioned by a variety of interacting factors, both biotic and abiotic, across different spatial and temporal scales.

The concept of liquid brain has recently gained prominence to describe a decentralized, flexible form of collective intelligence observed in certain organisms [4, 5]. These include ants and other eusocial insects, but also immune systems, slime molds, and some microbiomes. The units in a liquid brain operate as “leaky integrators” [6, 7], responding non-linearly to local encounter rates by integrating information and activity from nearby agents. Unlike traditional “solid brains” where connectivity is fixed, the network’s structure in a liquid brain is fluid. This fluidity arises from the agents’ movement, which continuously reshapes the patterns of interaction among them.

While liquid brains offer the potential for understanding generalizable properties across several living systems exhibiting collective intelligence, this framework must account for essential ecological processes and scales often overlooked in complex systems approaches [3]. For instance, behavioral heterogeneity, both within and between individuals in a population, plays a significant role in shaping ecological dynamics across multiple levels of organization [8]. This heterogeneity is critical for adaptability and resilience in animal populations, as it can buffer them against environmental fluctuations [9]. It also influences movement and dispersal, affecting the colonization of new habitats and the navigation of fragmented landscapes [10, 11]. Equally fundamental are ecological trade-offs, such as those involving body size, reproductive effort, and foraging strategies. These trade-offs reflect the balancing act organisms face when allocating limited resources to competing needs, influencing behavior, individual fitness, population dynamics, community interactions, and ecosystem processes. For example, in foraging strategies, the exploitation-exploration trade-off refers to the balance organisms must strike between efficiently utilizing known resources (exploitation) and searching for new, potentially better resources (exploration). Optimizing this trade-off is critical for resource acquisition in dynamic and uncertain environments [12, 13]. Additionally, the concept of collective intelligence must consider the role of adaptive behavior, as well as the existence of fluctuating densities and transients (phases of seemingly stable but non-stationary behavior), which can ultimately lead to significant shifts in state [14]. This natural variability underscores the importance of flexibility and responsiveness in collective systems, which are essential for navigating ecological and environmental variability.

Here, we leverage a liquid brain framework to investigate the mechanisms underlying collective foraging efficiency in ants. Building upon foundational mathematical models (e.g. [15, 16, 17]), our approach advances beyond theoretical constructs by validating the model with ecologically realistic data that includes detailed quantification of movement behavior. This method contrasts with precedent studies that often relied on scattered empirical datasets limited by scale and computational constraints. To conceptualize the liquid brain model, we examined the foraging dynamics of *Aphaenogaster senilis*, using an empirical setup that emulates the spatial and temporal scales of the foraging process in this ant species. *A. senilis* is characterized by inherently sparse networks of active foragers displaying heterogeneous movement patterns [18, 19, 20]. In these colonies, only a small proportion (10–50 individuals) actively engages in foraging, with the number of active agents fluctuating dynamically as individuals alternate between nest and foraging activity [21]. We incorporated these empirical insights, including ecologically realistic spatiotemporal scales, behavioral heterogeneity, foraging trade-offs, and time-adaptive strategies, to extend the liquid brain framework beyond its traditional conception as stationary, densely connected interacting networks. Using the extended liquid brain model, we investigated how *A. senilis* colonies leverage heterogeneous movement behaviors (encompassing both individual variability and socially guided motion) to manage sparse connectivity and enhance collective foraging efficiency across natural ecological scales.

## Materials and methods

### Study species

We conducted foraging experiments with *Aphaenogaster senilis*, a Mediterranean ant species known for its flexible foraging strategies, ranging from solitary scavenging to group recruitment [22]. Their colonies consist of hundreds of monomorphic workers (i.e. with no morphological casts), ranging from 100 to 3000 [23], but only about 10% to 20% of them leave the nest to perform tasks in the open space. They forage under a wide range of temperatures during spring and summer [22], and exhibit an omnivorous diet [24]. *A. senilis* usually forage within a 2-meter distance from the nest [25], which is about the scales used in our laboratory experiments, and are observed to display heterogeneous spatial distribution [18, 19] and movement behavior [20]. Overall, this species shows an interesting display of behavioral plasticity, which makes it a suitable model to study the questions presented in this paper.

### Animal holding

A colony of *Aphaenogaster senilis* (including workers, queens and brood) was collected at the university campus of Universitat Autónoma de Barcelona, Cerdanyola (Barcelona) in mid-April 2018. The colony was transported by car the same day (1-hour trip) to the laboratory in the Center for Advanced Studies of Blanes (CEAB). Upon arrival, the animals were kept under constant temperature (27°C) and a 14:10 light cycle. The colony was allocated to an artificial nest consisting of an opaquely covered plastic box (11 *×* 8 *×* 3 cm). An opaque tube connected the nest to a similar uncovered plastic box with ventilation, for ants to get rid of debris and dead bodies. Outside of the foraging experiment periods, this setup was connected to the feeding arena (43 *×* 28 *×* 7 cm), where ants were fed the same amount of food and with the same frequency as in the experimental condition (see below). Water was provided *ad libitum* with a small plastic deposit with a piece of cotton connected to the nest, which was refilled regularly.

Ants were gently manipulated during colony extraction, transport, housing conditions, and experiments to ensure their well-being and best ethical practices.

### Foraging experiments

Foraging experiments with *A. senilis* took place twice a week, with two trials (i.e. foraging bouts of 3 hours) per day, one in the morning (from 10:00 to 13:00) and another one in the afternoon (from 14:00 to 17:00). During the experiments, a large honeycomb-shaped arena (2 *×* 1 m^2^, Figs. 4C, 5B) was connected to the nest, allowing the ants to freely forage. This Y-maze configuration is commonly used in studies of animal decision-making (e.g. [26, 27]), providing a reliable method for quantifying behavioral choices, such as movements to the left, right, or back. Furthermore, this structure is well-suited for analysis using network theory and models from statistical physics (e.g. [28, 21]).

**Table 1:** Liquid brain model parameter descriptions and values. The parameters are organized into three categories, (i) those related to the information transfer (Eq. 1), (ii) parameters governing ant movement behavior (i.e. movement heterogeneity and social copying, see below), and (iii) parameters associated with the Gillespie algorithm (see Temporal scale section below). A summary of the parameters and their initial values in the simulation are provided in the table. A detailed description of each parameter is provided throughout the Methods section. The information transfer parameters are as follows. (a) *S_i_*, level of activity of agent *i*, being *S_i_ <* 0 inactive, and active otherwise. (b) *g_i_*, sensitivity degree of agent *i*, with larger values implying a stronger influence from local neighbors activity. (c) *J_ij_*, coefficient matrix representing the semantics of information, where the pair of states of agents *i* and *j* modulates the integration of local activity (similarly to *g*). The states are related to whether the corresponding ant knows a food location (state = 1) or not (state = 0). (d) Θ, a threshold of minimal activity for ants to remain active. The Gillespie parameters are as follows. (a) *N*, number of foragers, either scouts or recruits, participating in the simulation. (b) *α*, rate of events in the nest, at which an ant leaves the nest or interacts with other ants. (c) *β*, rate of events in the arena, at which an ant moves, collects food, or interacts with other ants. (iv) *γ*, rate of spontaneous activation, signifying a probability of an inactive ant (*S_i_ <* 0) to become active (*S_i_* = *U* (0, 1)). The behavior parameters are the following. (a) *ρ*, the proportion of ants moving as scouts, being 1 *− ρ* the proportion of recruits. (b) *ɛ*, the proportion of ants that can be informed about food locations via social copying.

| Parameter | Description | Value |
| --- | --- | --- |
| $S$ | Activity level | 0 |
| $g_i$ | Sensitivity to local activity | $U(0, 1)$ |
| $J_{ij}$ | Information semantics | $\begin{pmatrix} J_{00} = 0.01 & J_{10} = 0.01 \\ J_{01} = 1.00 & J_{11} = 1.00 \end{pmatrix}$ |
| $\Theta$ | Decision threshold | $10^{-10}$ |
| $\rho$ | Proportion of scouts | 0.5 |
| $\epsilon$ | Proportion of social ants | 1 |
| $N$ | Number of agents | 100 |
| $\alpha$ | Rate of events in nest | $2 \cdot 10^{-3} \text{ s}^{-1}$ |
| $\beta$ | Rate of events in arena | $6 \cdot 10^{-1} \text{ s}^{-1}$ |
| $\gamma$ | Rate of spontaneous activation | $1 \cdot 10^{-5} \text{ s}^{-1}$ |

We considered two scenarios: an experimental condition with food resources in the arena (Food), and a control scenario without food (No-Food). The Food scenario emulated a highly predictable environment; food was consistently placed in the same location every trial and the arena was not cleaned after each foraging bout. Therefore, the environment was strongly predictable and ants were allowed to trace previous odors or learn about the disposition of the food items (see supplementary text). Food resources consisted of two *Tenebrio molitor* larvae cut into six pieces and distributed in the vertices of two hexagons of the arena, representing two food patches. The patches were equidistant from the nest (about 65 cm from the nest, Fig. 4,5), one oriented towards the left and the other to the right. In the No-Food, food was absent and the arena was cleaned with alcohol 70% after each foraging bout.

The experimental sequence was structured as follows. Initially, the colony underwent a 15-day starvation period to stimulate foraging activity. This was followed by a two-week conditioning phase, to ensure ants got familiar with the arena’s shape, size, and food distribution (as in the Food condition). We cleaned the arena with alcohol and subsequently, we conducted 10 experimental replicates under Food conditions. Afterward, a two-week acclimatization period ensued, where ants were allowed to explore a cleaned arena without food (No-Food). Finally, we conducted 12 replicates under No-Food conditions. Importantly, across all experimental phases (Food, No-Food, and acclimatization), the ants were provided with a consistent diet (two cut larvae of *Tenebrio molitor*, twice per week) ensuring uniform nutritional conditions throughout the study.

### The liquid brain model

We explored the potential of liquid brains to reproduce and understand the mechanisms underlying *Aphaenogaster senilis* foraging dynamics. The empirical data from the experiments was used to conceptualize, parameterize, and validate the model. The model incorporates three main components: (i) an information transmission process (Eq. 1), (ii) a set of movement and social behavioral rules, and (iii) a continuous representation of time implemented using the Gillespie algorithm [29]. A schematic representation of the model is provided in Fig. 1. Unlike previous studies [30, 31], our approach introduces the use of discrete space, which simplifies the characterization of movement behavior and information flows, while also incorporating natural scales that closely resemble the foraging spatiotemporal patterns of *A. senilis* in the field.

**Figure 1:**
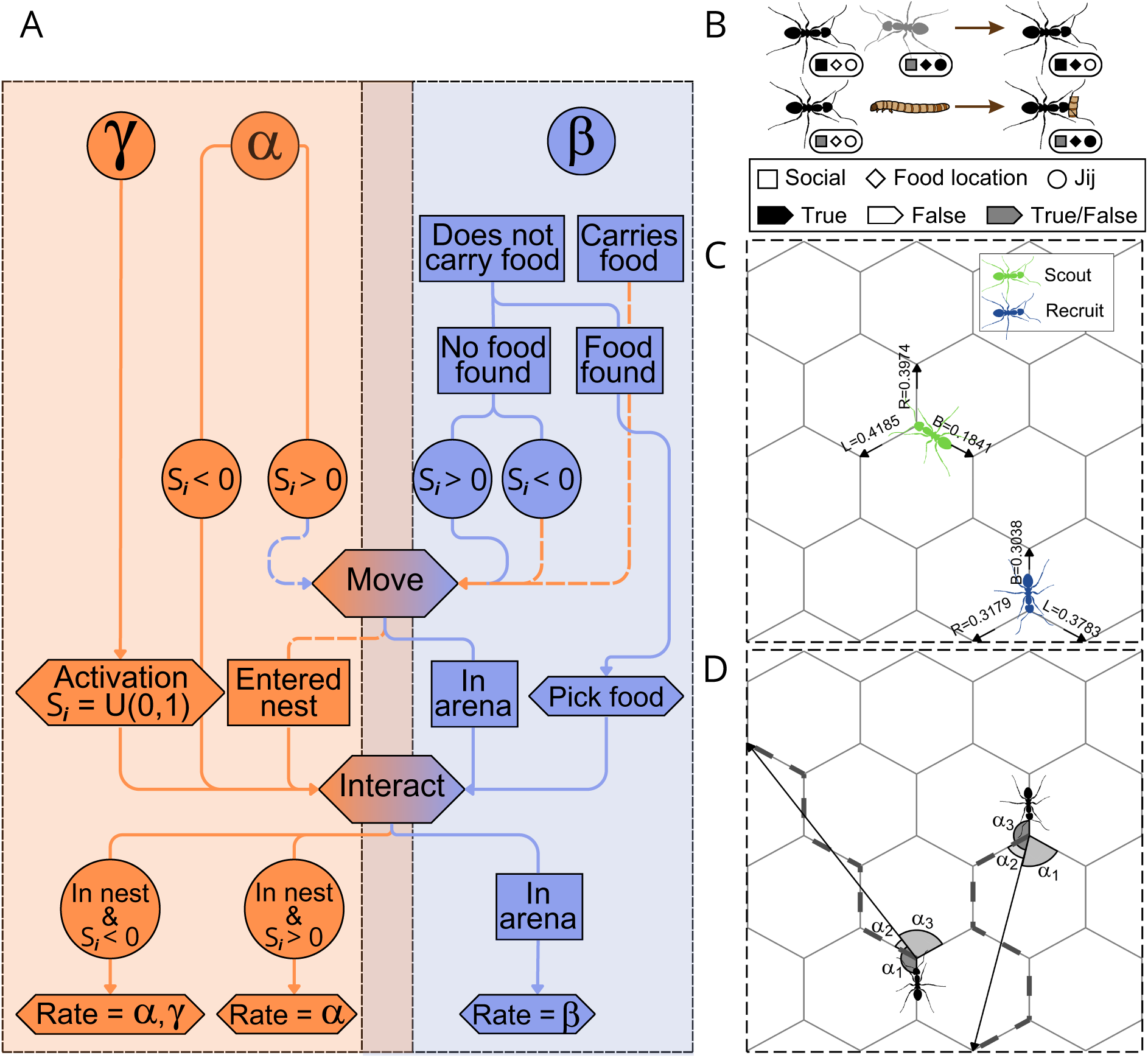
Illustrative flow chart of the model. A, the basic decision tree of the model in each iteration (see Table 1 for parameter details): an ant is selected from a pool in the nest (with a rate *α* or *γ*, orange-colored flow) or the arena (with a rate *β*, blue colored flow), the ant can perform some tasks if some condition is met, and finally it interacts with its neighbors. Conditional checks based on parameter values are depicted in the diagram with circles. The parameters *α*, *β* and *γ* respond to the rate at different ant actions occur, while parameter *S_i_* depicts the activity level of ant *i* (Eq. 1 and Table 1). Squares signify special conditional checks (e.g. food is found). Dashed lines illustrate transition states, in which the ant is going from the nest to the arena or vice-versa. Arrows connected to an hexagonal polygon imply an action (e.g. if food is found, it is then picked). Note at the end of each iteration in the model, a new rate is assigned to the selected ant. B, illustrative examples of interactions ant-ant (top) and ant-food (bottom). During interactions, ants adjust their activity levels (*S_i_*) and exchange information. A boolean (colors) vector of characteristics (symbols) describes whether an ant is social, knows about food locations, or has found food. Note that sociality is a behavioral trait of a particular ant that does not change as a result of an interaction (see Movement and sociality section below). Likewise, the outcome of some interactions is not altered by sociality (grey color). C, a portion of the lattice with ants displaying non-oriented movement, distinguishing the turning probabilities of scouts (green) and recruits (blue). D, a portion of the lattice with ants exhibiting oriented behavior, such as going back to the nest (homing) or being directed to a food location (through social copying). The turning probability depends on the angle (*α*) between the vector pointing to the target and the vector pointing to each possible direction (see Eqs. 2, 3).

#### Information transmission

Liquid brains model information transmission in a dynamic network of interactions. Agents move through space, interacting with one another and either amplifying or reducing the activity of others in a threshold-like manner, represented in Eq. 1[30, 31].

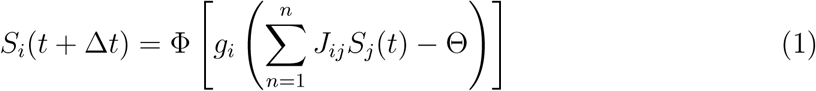

Agents’ activity (*S_i_*) determines their task allocation. An agent engages in foraging activities (exploring the lattice and collecting food) while *S_i_ >* 0. Otherwise, (*S_i_ <* 0) they will return to (or remain in) the nest. This activity level *S_i_* is influenced by the activity levels of local neighbors (agents *j*) that the focal agent *i* interacts with, considering up to *n* = 4 neighbors. The relationship between the neighbors’ activity and the focal agent’s response is governed by a sigmoid function (hyperbolic tangent, Φ), commonly used in neuronal models due to its threshold-like shape, zero-centered values, and range from −1 to 1. Note that for agent *i* to remain active (*S_i_ >* 0), the activity of its neighbors must exceed a minimal threshold (Θ).

Each agent has an intrinsic weight (*g_i_*), which represents its sensitivity to local information. As *g* increases, Φ approaches a step-function, making the agent’s activity (*S_i_*) more dependent on the neighbors’ integrated activity.

In addition, an extrinsic weight (*J_ij_*), which is universal across all agents, represents a set of states or rules that influence how information is processed. In the context of foraging, we considered these states to be related to food resources. Agents that find food (*state* = 1) become more influential than those that do not (*state* = 0). Practically, this means interactions with food-informed neighbors are more rewarding (i.e. *J*_01_ and *J*_11_, see Table 1) than with naive individuals who haven’t found food (i.e. *J*_00_ and *J*_10_).

#### Movement behavior and sociality

On top of the information transmission, we incorporated a social copying mechanism, that enables scouts to direct recruits to previously visited food locations. During an interaction with successful scouts, recruits are informed about the exact location of the last visited food source by the scout (Fig. 1B). We considered that informed ants can orient themselves towards food, having a preferred direction, but we are not describing the actual mechanism (recursive antennal contacts with informed ants, nestmate odour gradients, or pheromone gradients). Therefore, social copying follows a simple conditional rule: if any neighbor *j* has found food, agent *i* switches to oriented movement behavior (see below), heading toward the known food location. The proportion of foragers (scouts and recruits) that can exchange this spatial information is controlled by a sociality parameter (*ɛ*, Table 1). We assumed all ants to be social and able to be informed by this mechanism (*ɛ* = 1, Table 1), but we also investigate how varying this proportion influences foraging efficiency (see Results section, Fig. 6).

We defined (i) a non-oriented movement, capturing the heterogeneity observed in the search behavior of this ant species (i.e. scouts or recruits), and (ii) an oriented movement, emulating the ants’ ability to navigate back to known locations, such as a food location or the nest.

For non-oriented movement, we used empirically estimated turning probabilities, separately for scouts and recruits, capturing the observed behavioral differences between these two roles (Fig. 1C).

For oriented movement, we introduced directional persistence toward a target (Fig. 1D). Ants switched to this behavior when returning to the nest (i.e. when their activity dropped below 0, *S_i_ <* 0, or they found food) or when moving toward a previously discovered food source (i.e. as informed through the social copying mechanism described above).

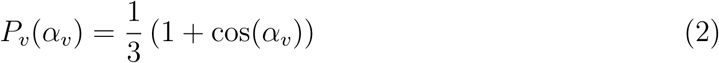

This directional persistence was modeled by Eq. 2, where the probability of moving to a given node (*P_v_*) depends on the angle (*α_v_*) between the vector pointing to the target and the vector pointing to the node. The probability of moving in the exact opposite direction of the target is zero (*P* (*π*) = 0).

At the edges of the arena, where only two vertices are available (due to the Y-maze shape), this probability was normalized (see Eq. 3).

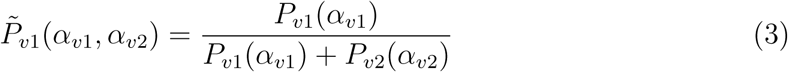

#### Temporal scale

To avoid using discrete simulation steps and capture the temporal dynamics of foraging, time was implemented in the model using Gillespie’s algorithm [29] (see also Table 1).

Briefly, the system is initialized at time 0 with a defined number of agents (*N* = 100). These agents can participate in different events within the nest (with rate *α* = 2*·*10*^−^*^3^ *s^−^*^1^) or in the foraging arena (with rate *β* = 6 *·* 10*^−^*^1^ *s^−^*^1^). Additionally, we included a small probability of ants becoming active in the nest without interaction (*γ* = 1 *·* 10*^−^*^5^ *s^−^*^1^). At each simulation step, a single event occurs, with its probability proportional to the number of agents available for that event (e.g. *P* (*α*) = *N_α_ · α/R*, where *R* is the sum of all event rates).

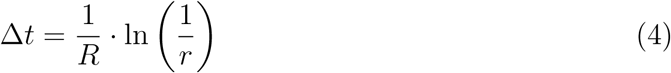

Time, measured in the same units as the event rates (i.e. seconds), is sampled from an exponential distribution. The characteristic time scale is determined by the sum of rates, *R*, multiplied by a random number, *r*, drawn from a uniform distribution bounded from 0 to 1(see Eq. 4). The simulations were run for 3 hours to match the temporal scales of the empirical data.

The values of the parameters *N*, *α*, and *β* were arbitrarily chosen, based on a qualitative inspection of the number of foragers, the rate at which foragers departed the nest, and their average speed while navigating the arena, respectively.

## Results

We conducted foraging experiments in which we allowed *A. senilis* colonies to explore a large, discrete, honeycomb-like arena (2 *×* 1 m^2^) for three hours (see Methods for details). This novel experimental setup features large spatiotemporal scales, closely reflecting field observations in this ant species. Additionally, it enables straight-forward quantification of decision-making behavior and provides a direct analogy to discrete lattices used in simulations. We considered an experimental condition with food (Food), where two food patches were consistently placed in the same locations across trials. These were compared to a control scenario with no food present (No-Food).

The foraging dynamics of ants showed a characteristic trend with two clear transition points (TP), as described in Cristin *et al* (2024) [21]. A few ants, generally from one to five, explored the arena until a food piece was spotted (exploration phase, TP1). This triggered the beginning of the exploitation phase, which lasted until all food was retrieved (TP2). Beyond this point, the system maintained momentum, with ants continuing to explore at peak activity levels for an additional 20–30 minutes before gradually returning to the nest.

### Scouts and recruits: behavioral and movement heterogeneity

We analyzed the search behavior of ants during the exploration phase. In the Food experimental condition, we focused on ant trajectories occurring before food discovery (the first 15 minutes on average). The No-Food control condition served as a null model for search movement, presuming that no additional processes (i.e. recruitment and patch exploitation) occurred.

Each ant trajectory was composed of a set of spatial coordinates, for which we calculated the distance from the nest and then averaged the result. The distribution of average distances from the nest exhibited two characteristic distance-to-nest scales, indicating a clear spatial organization within the arena (Fig. 2). These two regimes, near and far from the nest, were separated by a breakpoint at 20 cm in the Food condition and 35 cm in the No-Food condition.

**Figure 2:**
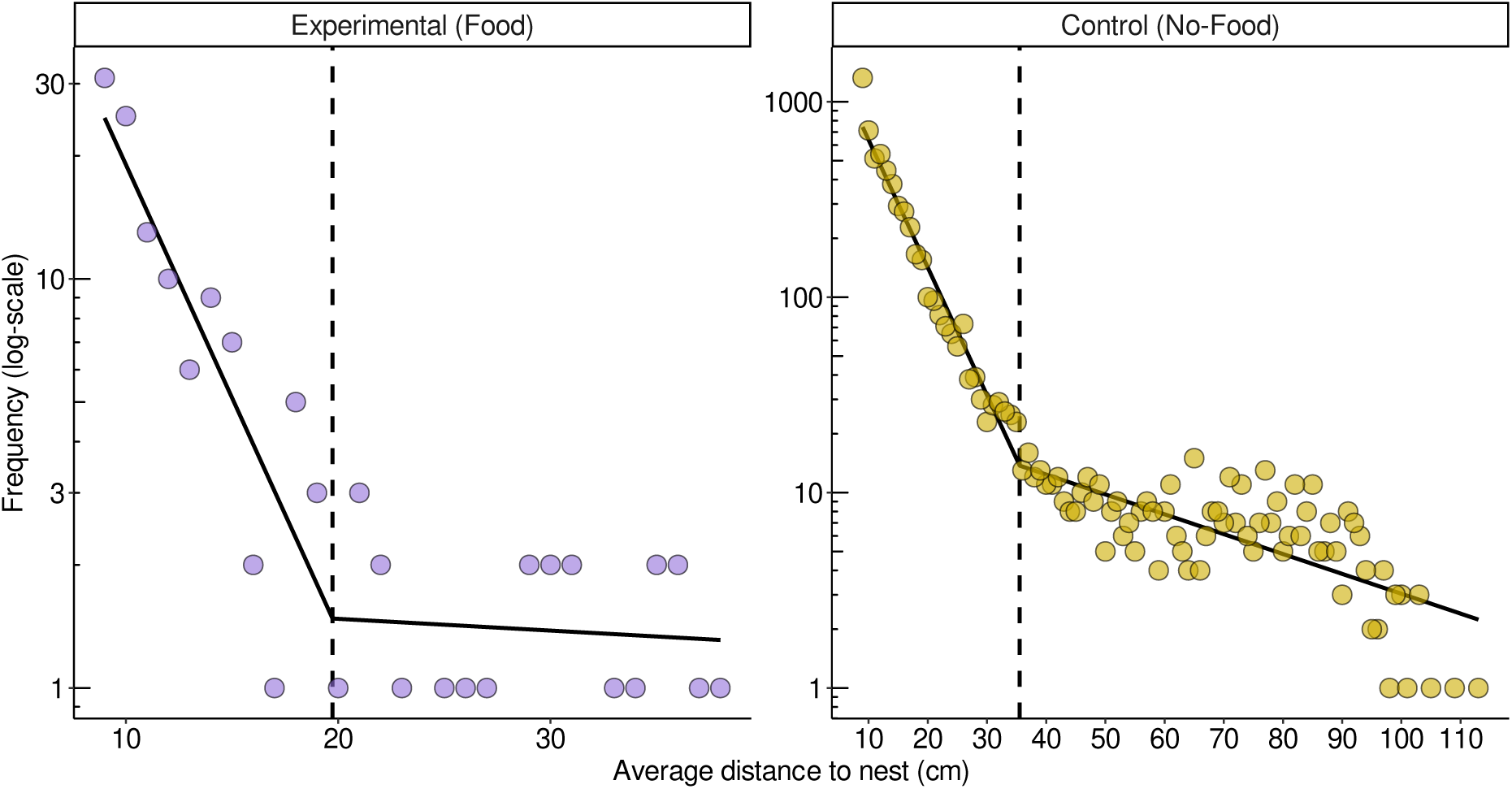
Distribution of average distances to the nest for different ant tracks during the exploration phase. We calculated the average distance-to-nest of each ant trajectory, accounting for all the trajectories before food was found (exploration phase). We show the distribution (frequency in log-scale, y-axis) of these distances (x-axis) of individual tracks (dots) for the Food (left panel) and No-Food (right panel) scenarios. A piecewise model was fitted on log-linear scales to depict the occurrence of two characteristic spatial scales. The dashed line splits these two regimes of nearby and far distances to the nest, around 20 and 35 cm for the Food and No-Food conditions, respectively. Note that the exploration phase lasts for the whole trial in the No-Food scenario (right panel), given the absence of food. Therefore, there is much more data available in this case than in the Food condition (left panel).

We propose that this spatial organization reflects the presence of two distinct roles observed in this ant species: scouts, which are exploratory ants venturing far from the nest, and recruits, which exhibit a marked preference for aggregating close to the nest, awaiting instructions from the scouts. Accordingly, we considered trajectories exceeding the breakpoint average distance to be scouts, or recruits otherwise. The breakpoint distance was specific to each condition, as explained above (see Fig. 2).

Computing the likelihood (i.e. probability density function) of encountering scouts and recruits at different distances from the nest further evidenced the differences in these two behaviors (Fig. 3). Recruits were mainly found within 20 cm from the nest (note the modes of the density in Fig. 3), occasionally exploring beyond 35 cm. Conversely, scouts ventured much farther, often exceeding distances of 60 cm from the nest, and displayed a more uniform distribution across intermediate and distant ranges from the nest. This uniformity in spatial distribution is more pronounced in the No-Food condition compared to the Food condition (see also supplementary text).

**Figure 3:**
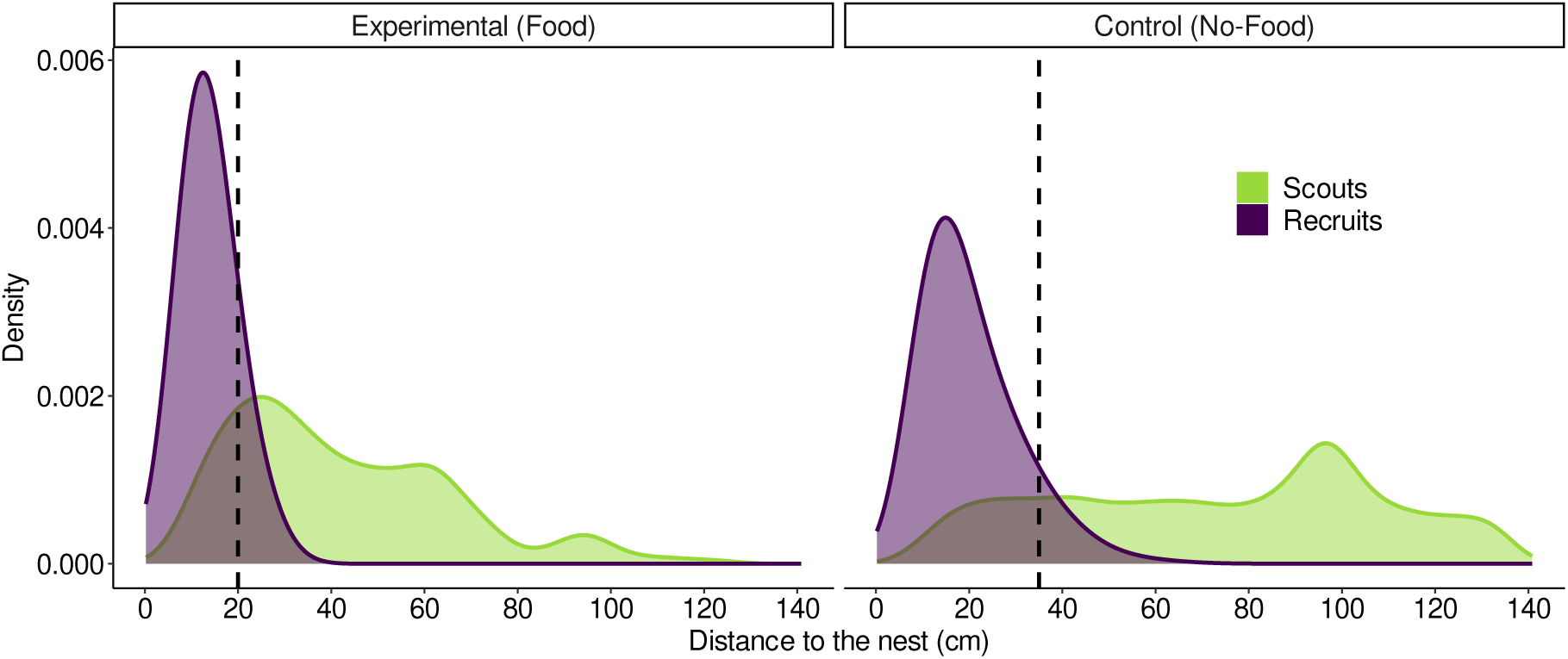
Heterogeneous spatial distribution of scouts and recruits. We show the density (y-axis) of scouts (green) and recruits (blue) across increasing distances to the nest (x-axis). This represents the likelihood of encountering a scout (or recruit) at a given distance from the nest. The dashed vertical lines signify the breakpoint used to classify ant trajectories into scouts and recruits (as shown in fig. 2). Note that recruits are mainly concentrated close to the nest (less than 20 cm), rarely exploring further than 35 cm away from it, while scouts cover a broad range of distances.

The arena setup played a key role in defining the spatial scales observed in each condition. In the Food condition, exploration time was limited by the moment of food discovery, after which social feedback and recruitment were triggered. Similarly, the spatial extent of exploration was bounded by the location of the food source, effectively reducing the likelihood of reaching far distances (see also supplementary text). Conversely, the spatial scale in the No-Food condition was solely constrained by the size of the arena (Fig. 3).

On top of this heterogeneous spatial distribution, the movement properties of the two types of ants were remarkably different from one another (Fig. 4). Recruits displayed lower velocities (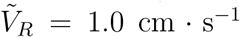, 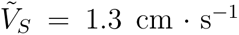, *p*-value *≈* 0) and directional persistence (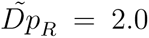 steps, 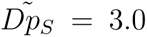 steps, *p*-value *≈* 0) than scouts (Fig. 4A, top panels). Moreover, recruits tended to enter and leave the nest with much higher frequency, resulting in covering less distance (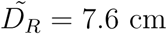, 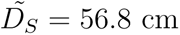, *p*-value *≈* 0) and less time (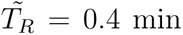, 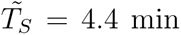, *p*-value *≈* 0) exploring (Fig. 4A, bottom panels).

**Figure 4:**
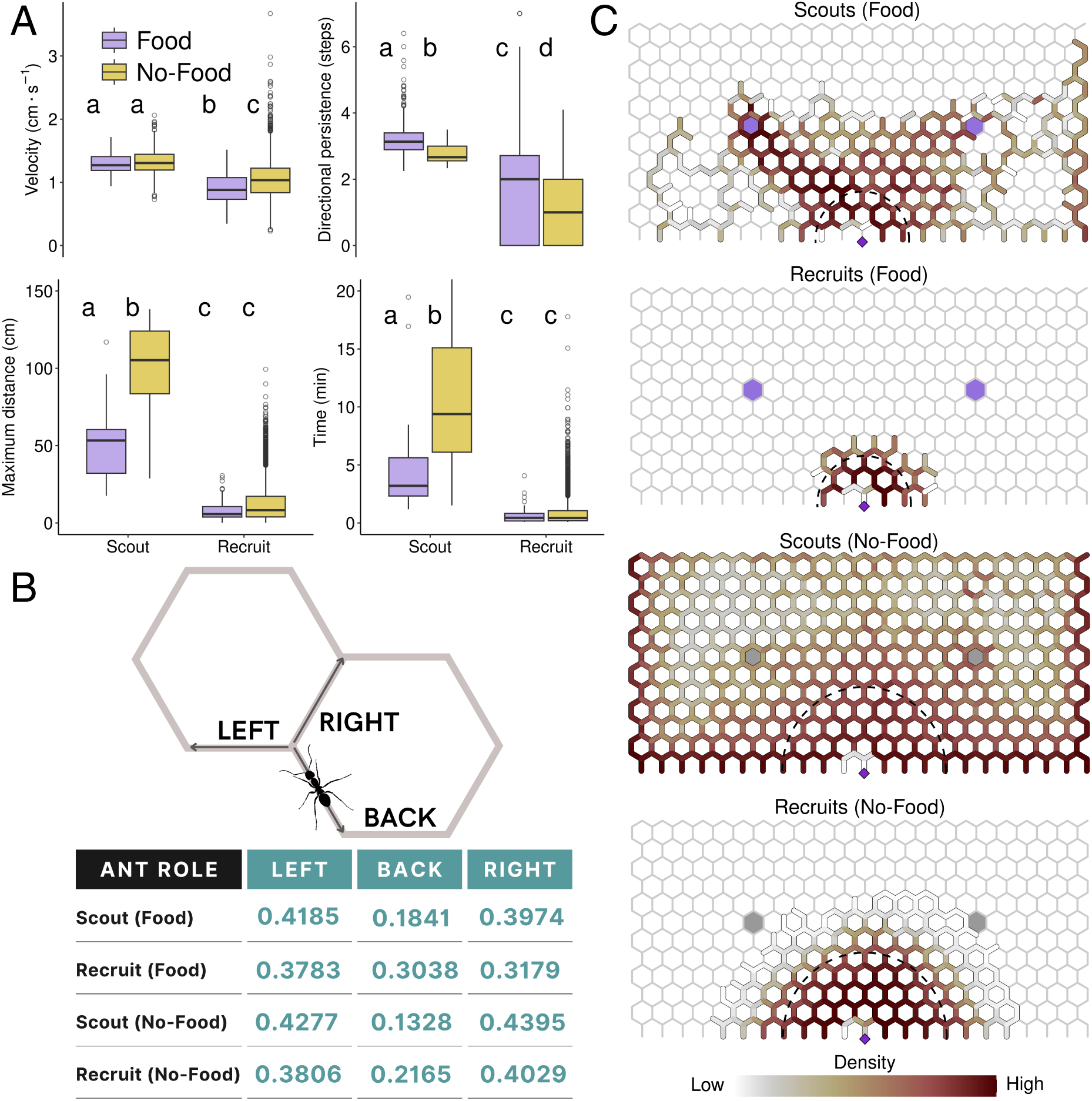
Movement heterogeneity of *A.senilis*. We characterized the movement of scouts and recruits with food (Food) and without food (No-Food) conditions. A, movement characterization based on ant trajectories: velocity (in cm *·* s*^−^*^1^), directional persistence (number of steps following left-right, zig-zagging turns), distance traveled (cumulative distance in cm), and time spent in the arena (in minutes). Letters identify groups belonging to the same distribution, according to the results of paired Wilcoxon tests (*α* = 0.99) using Bonferroni correction. B, turning behavior, signifying the probability of a right, back, or left turns in the trajectory. C, average density, measured as the number of positions detected in each segment delimited by two adjacent nodes. The purple diamond shows the entry point to the arena (nest), and the purple hexagons depict the position of food in the Food condition (also shown in grey for the No-Food scenario as a visual reference). The dashed semicircle depicts the threshold distance that separates the two characteristic scales of observed distances from the nest, used to classify scouts and recruits (see Fig. 2). Heat maps show a qualitative scale to aid comparative visualization among panels.

The combination of these microscopic movement properties, including velocity, directional persistence, and turns (Fig. 4A top panels and 4B), directly influenced the overall time spent, distance traveled, and space utilized within the arena (Fig. 4C). For example, scouts made fewer backward turns compared to recruits. This pattern emerged from a higher frequency of ‘left-right’ (zigzagging) turning sequences that, on the honeycomb lattice, resulted in greater directional persistence, leading to straighter trajectories and larger distances covered. In contrast, recruits exhibited nearly equal probabilities of turning left, right, or backward, resulting in diffusive-like trajectories. These movements were often restricted to areas near the nest, as shown in figure 4C.

The observed distinct behaviors of scouts and recruits were consistent across the Food and No-Food conditions, though some subtle differences appeared. In the absence of food resources, both scouts and recruits exhibited lower probabilities of making backward turns compared to when food was present (Fig. 4B), but showed less directional persistence (zig-zagging), and were slightly faster (Fig. 4A). These slight differences in microscopic movement rules when comparing arenas with and without food, could not explain by themselves the differences observed in macroscopic space use patterns (Figs. 2 and 3). For instance, although scouts exhibited similar velocities and turns (Fig. 4A,B), their macroscopic exploratory patterns differed drastically between arenas with and without food (Fig. 4C). In the Food condition, scout paths closely followed optimal routes (minimal Euclidean distances) that connected the nest to the two food patches. This likely occurred because the arenas were not cleaned between trials, allowing odor cues to persist and retain information about food locations. In contrast, in the No-Food condition cues were erased between trials, and scouts displayed trajectories radiating from the nest in mostly straight lines toward the arena’s periphery, resulting in paths that consistently reached the arena’s borders. These clear differences between optimized (with food) and non-optimized (without food) space use patterns were further confirmed by measuring scouts’ spreading properties from the nest, as reflected in first-passage times (supplementary text, fig. S1).

### Foraging ants as a liquid brain

Foraging in *A. senilis* colonies is a dynamic, non-stationary process comprising three distinct phases (Fig. 5A), delineated by two key transition points (TP). The first transition point (TP1) marks the discovery of the initial food item, while the second (TP2) occurs when the last food item is retrieved. Before food is detected, a small number of scouts (typically fewer than five) search the arena, while most recruits remain near the nest. Upon food discovery (TP1), scouts transmit information to recruits, and through a social feedback mechanism known as group recruitment, a surge of up to 40 recruits end up leaving the nest over approximately 30 minutes. During group recruitment, scouts and recruits form back-and-forth trails connecting the nest to food patches. After all the food is collected (TP2), the system slows down, leaving only a few ants active in the arena.

**Figure 5:**
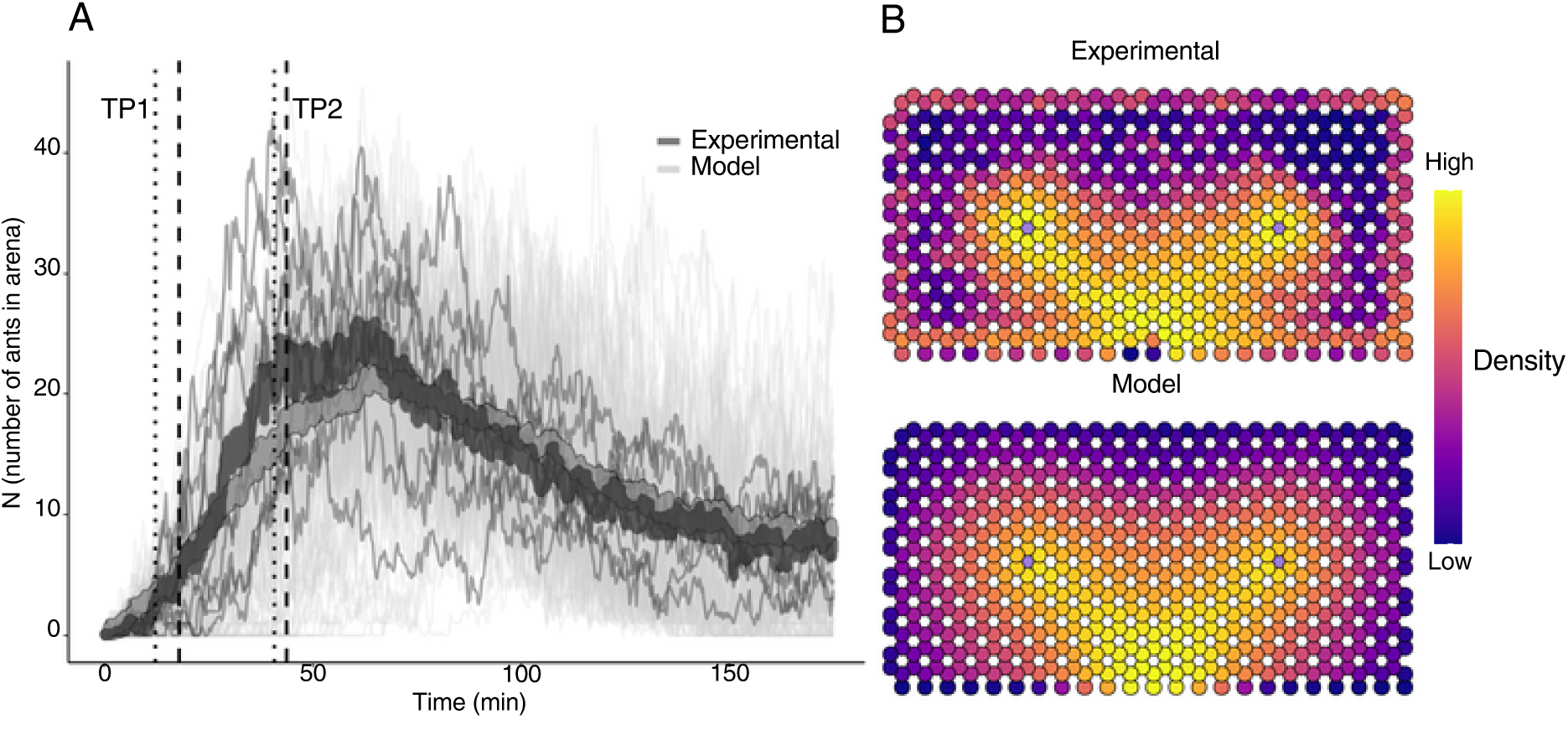
Foraging dynamics of *A. senilis*: comparison of 10 experimental (Food) foraging trials with 100 simulated stochastic replicates. We carried out 100 simulations with the liquid brain model (see Table 1 in Methods) replicating the same food placements and spatial scales as in the experimental conditions. A, comparison of population dynamics (number of ants in the arena, N) between the experiments with food (dark grey) and simulations (light grey). The thick curves represent the average, whilst the thin lines represent each experiment or simulation. The vertical lines signify two key transition points (TP) in the foraging process: (TP1) the first time food is found and (TP2) the time at which all food has been collected. Dashed lines show the mean of these times for the simulations, whilst dotted lines show the mean of the experimental data. B, average density (number of ants in each node) heatmap for the experimental data (Experimental, top panel) and the simulations (Model, bottom panel).

Our liquid brain model incorporates two fundamental components that enable a quantitative replication of foraging dynamics across both space and time (Fig. 5). The first component is the modulation of individual ant activity as a leaky integrator [7]. Each ant integrates the activity levels and food-related information from neighboring ants, responding to local encounter rates through a sigmoid function, analogous to neuronal responses. The second component is the liquid nature of network connectivity, which emerges from the ants’ movement patterns. These patterns include non-oriented movement, distinguishing scout and recruit behaviors (Fig. 4B), and oriented movement towards the nest or previously discovered food resources. Ants are assumed to exhibit oriented movement toward the nest (homing behavior) after finding food or after an unsuccessful foraging bout (i.e. when their activation levels drop below a critical threshold). On the other hand, oriented movement toward food sources is socially driven and occurs when successful scouts inform recruits about food locations.

Our model assumes an initial composition of 50% scouts and 50% recruits, although substantial variability in these proportions was observed across foraging trials (supplementary text, fig. S2). This variability underscores the flexible, adaptive nature of role allocation within *A. senilis* colonies and highlights the importance of incorporating such heterogeneity into the liquid brain framework.

Using our model (see Fig. 1, Eq. 1 and Table 1 in Methods), we quantitatively reproduced the empirically observed dynamics of *A. senilis* foraging described above. Once the first food item is found (exploration phase, first dashed line in Fig. 5A) information is rapidly transmitted throughout the colony, creating positive feedback from which self-organized recruitment emerges (exploitation phase). This amplification in the activity of ants arises from an increase in their excitability (or sensitivity to information), encoded in the matrix *J_ij_*. Accordingly, as ants interact with successful foragers, whether in the nest or along the pathways connecting with food patches, their activity is amplified non-linearly, continuously reinforcing recruitment. In this process, some ants inform their nestmates about the food locations, further contributing to forming trails toward food patches. After both food patches have been depleted (second dashed line in Fig. 5A), the overall activity progressively declines as interactions with successful foragers become less frequent, ultimately resulting in ants returning to the nest. Therefore, both ant recruitment and withdrawal to the nest are self-regulated. These behaviors emerged from the synchronized activity of the foragers, shaped by their movements, local encounter rates, and information transmission. Notably, the movement behavior of scouts or recruits affected search efficiency but did not govern population dynamics, which is primarily driven by social feedback mechanisms during the exploitation phase (supplementary text, fig. S3).

In Fig. 5B, we present the average spatial occupancy patterns of ants observed in experiments and simulations using the model. Both the empirical data and the simulations show that ant traffic is predominantly concentrated in three key areas: (1) the nest, (2) the food patches, and (3) the pathways connecting the nest to the food sources. However, a notable discrepancy emerges: nodes on the periphery of the arena exhibit significant occupancy in the experiments but not in the simulations (Fig. 4C). This highlights the behavior of ants that avoid patch exploitation and social feedback, instead continuing to explore far from the nest despite the presence of established resources.

### Foraging efficiency and the exploitation-exploration trade-off

We investigated how movement behavior shapes the foraging patterns of *A. senilis* colonies (Fig. 6), focusing on the interplay between movement heterogeneity (i.e. the proportion of scouts and recruits) and socially oriented movement (i.e. the extent of social copying). Specifically, we analyzed how these two factors influence foraging efficiency across two distinct phases of the process. In the exploration phase (Fig. 6A), we measured search efficiency as the time required to find the first food item. In the subsequent exploitation phase (Fig. 6B), we measured the exploitation efficiency as the time required to collect all food items present in the arena. The sum of these two metrics defines the total foraging efficiency (Fig. 6C), showing how ant colonies balance the two processes (exploration or exploitation).

**Figure 6:**
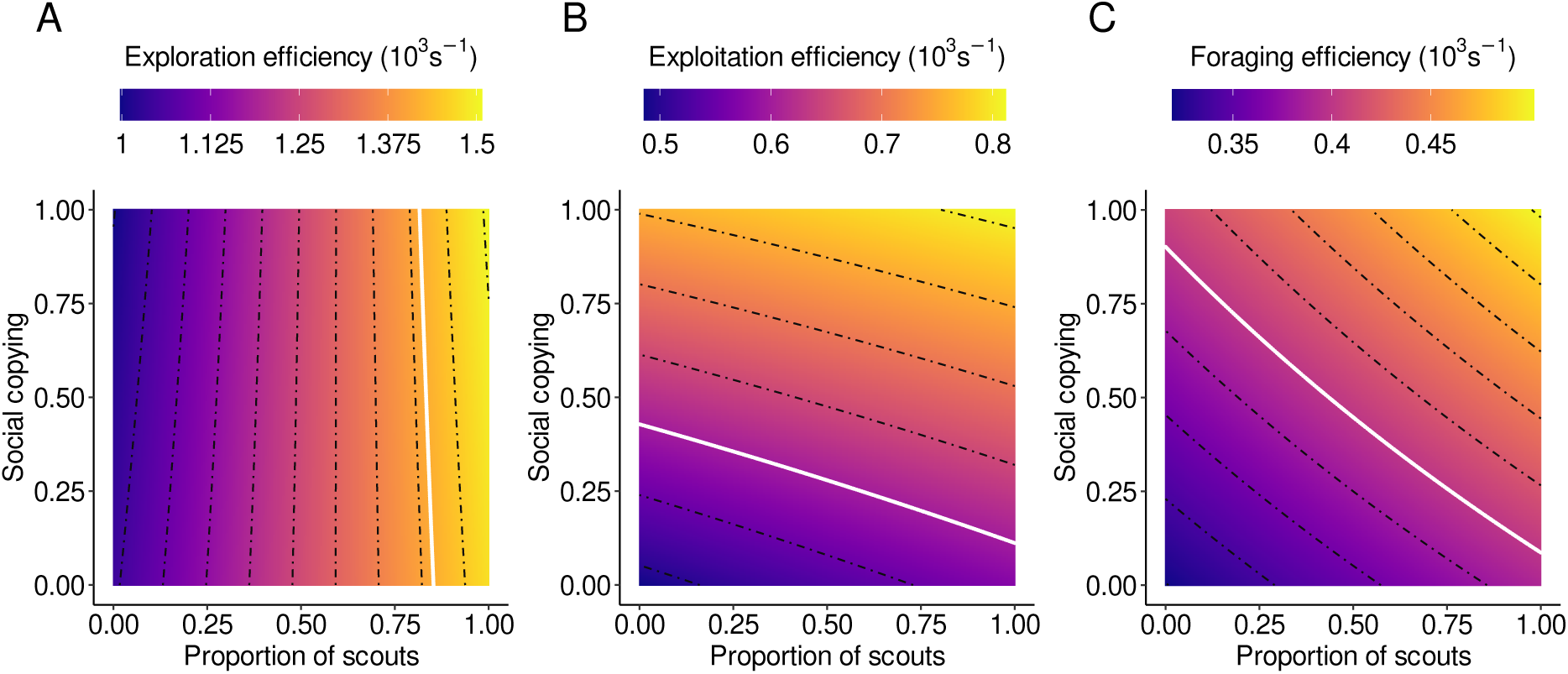
Behavioral heterogeneity and sociality as drivers of *A. senilis* foraging efficiency.. We represented movement heterogeneity (x-axis) as the proportion of scouts, and social copying (y-axis) as the proportion of ants that can be informed about the location of food targets. The metrics represented in the figure are expressed as rates (in kilo seconds*^−^*^1^, to ease the legend’s visualization), indicating the frequency at which foraging events occur. Note that each pixel in the landscape corresponds to the median efficiency computed throughout 100 simulations. The colors, deep blue to yellow, signify a gradient of increasing efficiency. The dotted black lines represent the contour isolines. The white lines represent the median empirical efficiency. A, exploration efficiency, the time span between the first ant leaving the nest and the discovery of the first food item, 1*/*(TP1 *−* min(t)). B, exploitation efficiency, the time spent collecting food after the first food item has been discovered, 1*/*(TP2 *−* TP1). C, foraging efficiency, the time required to find and collect all food, 1*/*(TP1 *−* min(t) + TP2).

We found exploration efficiency to be directly proportional to the initial relative number of scouts (Fig. 6A). Scouts were more efficient searchers than recruits, decreasing first-passage times by covering larger distances in a lower amount of time (supplementary text, fig. S4). The gradient in Fig. 6A showed that increasing the proportion of scouts decreased the time required to discover food. It was expected that social copying (y-axis) did not influence exploration efficiency, since it did not come into play during the exploration phase. Notably, the exploration efficiency observed in the empirical data (white line) was confined to a narrow region of the parameter space, where the proportion of scouts exceeds 75%. However, our Food trials revealed considerable variability in the initial relative number of scouts. Despite this variability, a high proportion of scouts among the ants leaving the nest was consistently necessary for efficient food discovery, irrespective of the level of social copying. Nonetheless, having some recruits present in the arena appeared also to play a crucial complementary role, likely by facilitating the rapid dissemination of information about food discoveries made by scouts. This initial presence of recruits may enhance the efficiency of transitioning from exploration to exploitation, ensuring that food resources are quickly and effectively utilized once discovered. Importantly, this feature is not well-captured by assessing first detection times as in (Fig. 6A). This balance between scout-driven exploration and recruit-mediated communication underscores the importance of movement heterogeneity in the exploration phase.

Exploitation efficiency improved as more ants were recruited and actively engaged in collecting food from patches. Therefore, resource exploitation was primarily driven by social copying. Interestingly, exploitation efficiency also increased with a higher proportion of scouts, as indicated by the diagonal gradient in the heatmap (Fig. 6B). It is noteworthy that in the experimental trials, some ants were observed exploring the periphery of the arena during the exploitation phase, highlighting the idea of simultaneous exploratory and exploitative behaviors (Fig. 5B). This suggests that both recruitment and a sufficient proportion of scouts are essential for optimizing exploitation processes. The proportion of scouts influenced the exploitation phase by increasing the likelihood of discovering new food patches while others are still being exploited, thereby enabling simultaneous and more synchronized exploitation of multiple patches (supplementary text, fig. S5).

When assessing overall foraging efficiency, encompassing both the exploration and exploitation phases, we observed a pronounced diagonal and symmetrical gradient (Fig. 6C). This pattern suggests that foraging efficiency improved similarly with increases in either scouts or socially-driven movement. At our experimental scales (i.e. spatial arrangement of food targets and arena size), scout proportion and social copying contributed additively to foraging success, displaying a clear compensatory role.

## Discussion

We quantitatively replicated *A. senilis* foraging patterns at ecologically realistic scales with a model describing a set of moving agents that follow neuronal-like interactions. Our empirical setup was specially designed to closely align with the foraging spatiotemporal scales of *A. senilis*. The foraging arena, featuring a honey-comb pattern, enabled precise quantification of simple behavioral choices (i.e. directional turns) and seamlessly integrated into our modeling framework. This approach allowed for a detailed analysis of ant movement, which we incorporated into a liquid brain model. Our results show that recruitment emerges from a feedback mechanism, driven by information transfer. This feedback is further modulated in the foraging area, where recruits and scouts play a key role in balancing exploration and exploitation processes, ultimately contributing to maximizing foraging efficiency.

Our results illustrate the connectivity of ant colonies as a dynamic, fluid property, with both the number of ants and the types of interactions evolving over time. Overall, in ‘low-dense’ liquid brains with few or spatially sparse agents, movement heterogeneity (i.e. scouts *vs.* recruits, oriented *vs.* non-oriented motion) plays a crucial role in sustaining and modulating the interaction network among ants. Individual heterogeneity within social groups manifests in various forms, such as differences in sex, age, or leadership roles [32, 33]. In many ant species, it is widely accepted to classify workers into two types: scouts and recruits. In our study, the distinction between scouts and recruits emerged from movement patterns related to differences in spatial occupancy and persistence within the arena [20, 34]. As ants discover food, their movements become more oriented. Food discoveries trigger social copying interactions, in which ants share information and guide others toward food sources. This social copying behavior feedback increases the number of ants actively participating in foraging and facilitates the transfer of resource information across the colony [7, 35, 36, 37]. Thus, the colony dynamics and foraging strategy are governed by a key feedback loop between movement patterns and information transfer. This loop ensures that ants remain engaged in tasks related to both resource exploration (via non-oriented motion) and exploitation (via oriented motion). Oriented motion, in particular, plays a critical role in creating dynamic bridges that connect the nest to food patches. As food sources become depleted, there is a general decrease in activity levels, compelling the ants to return to the nest.

Our findings reveal that exploration is primarily driven by the non-oriented movement of scouts, while resource exploitation is largely facilitated by oriented movement through social copying. However, this distinction shows only part of the story: recruits also play a critical role during the exploration phase, while scouts can take on a prominent role during the exploitation phase.

During exploration, we observed that increasing the number of scouts significantly reduced the time required to locate food, thereby improving exploration efficiency. Because scouts maximize dispersal speed and spatial coverage, one would expect exploration to rely solely on scouts. However, this assumption does not fully capture the dynamics of ant foraging, where information transfer among scouts and recruits plays a pivotal role in enhancing collective success. This is reflected in our empirical observations, wherein a proportion of recruits consistently departed alongside scouts at the onset of foraging bouts. The initial ratio of scouts to recruits varied significantly across experimental trials and is likely to differ among colonies as well [38]. The presence of recruits near the nest increases local densities, creating communication hubs during the exploration phase. These hubs may facilitate enhanced information transfer and enable the immediate exploitation of food sources upon discovery.

In environments with high food predictability, scouts optimized space occupancy thereby reducing exploration times [21]. The colony adjusted space occupancy in response to food presence, even though the microscopic movement rules (turning angles) and number of scouts remained similar to the control (No-Food) conditions. These results suggest that scouts likely utilized olfactory cues [39] or spatial memory [40] to navigate efficiently toward anticipated food sources. Indeed, different kinds of memory help improve foraging efficiency in temporally predictable environments [41]. The learning behavior observed in scouts, parallels observations in other social organisms. For instance, honeybees navigating predictable floral landscapes [42, 43], seabirds exploiting consistent food subsidies [44], flocking birds following established migratory routes [45], and wolves hunting within familiar territories [46] all demonstrate how the ability to anticipate resource locations can significantly enhance collective foraging efficiency.

During exploitation, social copying behavior prompts ants to focus on exploiting a single resource patch before moving to another. This emerges from an information-driven competition compelling ants to exploit food patches with different recruitment efforts, even when their quality is similar [47]. This is indeed a widespread phenomenon across several animal taxa: a minimal level of social copying provides a critical advantage for efficiently utilizing resource patches [41, 48, 49]. These density-dependent feedback mechanisms foster coordinated exploitation and positively influence collective foraging dynamics [50, 51]. Notwithstanding, our results suggest this same behavior can hinder the simultaneous exploitation of multiple patches. For ants featuring similar levels of social copying, we found that higher proportions of scouts enhanced exploitation efficiency by increasing the probability of discovering alternative food sources. In addition to a probabilistic decision to follow information cues [15], our results suggest that heterogeneous movement alone provides a mechanism to regulate the connectivity of the information network, thereby modulating the feedback generated by social copying.

In general, increasing the number of foragers improves food discovery in collective systems [52, 53]. However, our findings demonstrate that the roles of scouts and recruits contribute differently to foraging efficiency, depending on the stage of the foraging process and modulated by both individual and social learning. At the observed scales, timings of food detection and gathering remained consistent across different proportions of scouts and varying levels of social copying. This result highlights the existence of multiple collective solutions capable of achieving comparable foraging efficiency. This coordinated variability modulates ant network connectivity and enables a balance between exploration and exploitation, providing resilience against errors and inefficiencies under uncertain conditions.

We showed that the liquid brain metaphor in conjunction with solid empirical data, provides a compelling framework for understanding the ecological mechanisms at play in multi-agent social systems. Behavioral heterogeneity serves as a key driver of population plasticity, enhancing collective efficiency (reviewed in [8]). We found a simple feedback mechanism at the core of self-emergent ant recruitment, operating without the need for additional cues (e.g. pheromone). This process was further modulated by distinct movement types that shaped connectivity patterns, simultaneously forming exploratory routes and interaction hubs (e.g. near the nest) where resource discoveries are efficiently transmitted. These mechanisms may reflect characteristic organizational principles of other liquid brains, such as T-cell recruitment in lymphatic nodes [54]. Further studies should address the extent of this plasticity. It remains unclear whether the proportions of scouts and recruits change over time, or whether individuals consistently maintain their roles (i.e. as ‘personalities’). Task switching has been proposed to be modulated through local and context-dependent interaction rates [7], potentially introducing an additional layer of plasticity and resilience to liquid cognitive systems. Altogether, feedback mechanisms are regulated by movement-driven reconfigurations of the information fluid network. These reconfigurations, in turn, foster an adaptive and robust form of collective intelligence, enabling the optimization of key collective processes such as foraging in ants. Analyzing the dynamics of specific mechanisms may uncover generalizable properties of self-organized systems [3, 55], where coordination emerges from local interactions without centralized control [1, 2, 56]. Accordingly, the processes and mechanisms we describe may be shared with other liquid brains, whether found in social organisms, such as ant colonies and human societies, or in decentralized biological or artificial systems that process information collectively.

## Acknowledgments

We would like to acknowledge Xavier Espadaler and Daniel Campos for vivid discussions on ant behavior and modeling. We also would like to express our deepest gratitude to Daniel Guerrero for his companionship and contributions to this research. His dedication, passion, talent, and insights were valuable in shaping this research. His passing meant a great loss to the team, but his influence will hopefully be forever reflected in this paper.

