## Supplementary_Material for "Foraging Ants as Liquid Brains: Movement Heterogeneity Shapes Collective Efficiency"

### Effects of resource predictability and group recruitment on scouts movement behavior

While we did not observe microscopic differences in the movement of scouts between the Food and No-Food conditions (Fig. 4B), we observed distinct macroscopic patterns. These differences were evident in metrics such as first-passage times (FPT) as a function of distance from the nest (fig. S1).

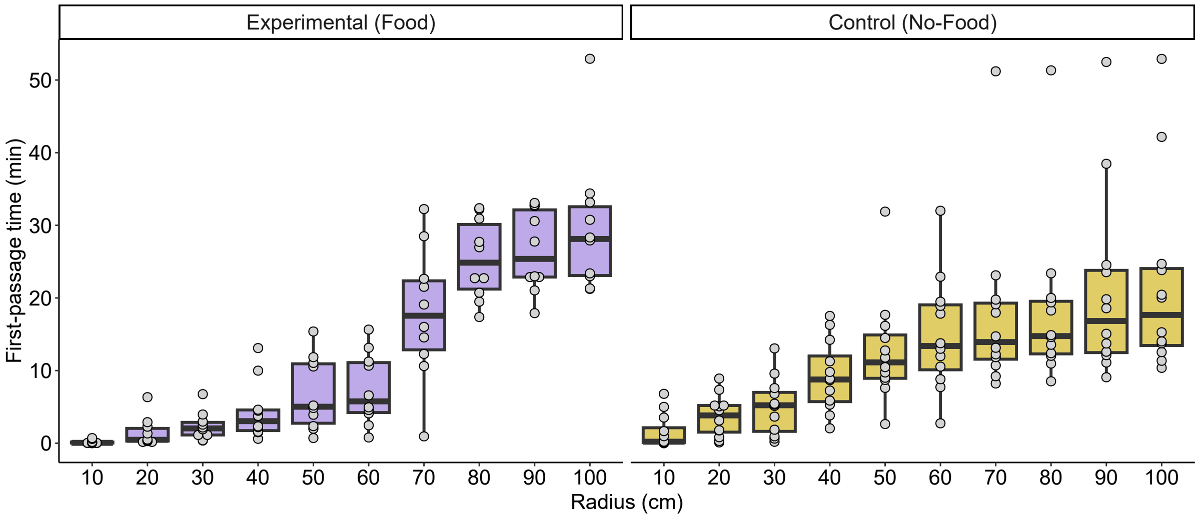


**Figure S1: Empirical first-passage times as a function of the distance from the nest.** We measured the time (y-axis) required for the colony to reach a certain distance from the nest (x-axis). We show the first-passage time distributions for the Food (left) and No-Food (right) conditions, with each dot signifying one of the 10 (12) Food (No-Food) replicates.

In both cases Food and No-Food conditions, the FPTs grew non linearly with distance. However, the curves are different. In the Food condition, the FPT increased with distance following a sigmoid curve, whereas in the No-Food condition, it followed an asymptotic curve. In addition, in the Food condition, scouts traversed through distances up to 60 cm considerably faster (3-fold on average, ranging from 1.9 to 8.8 fold) than in the No-Food scenario. On the contrary, beyond 65 cm far distances were reached about 60% slower in Food compared to No-Food conditions. Notably, food patches were located at 65 cm distance from the nest.

The observed differences may be attributed to the scouts' memory, sensory abilities, and behavioral priorities. In the Food trials, food patches were consistently located in the same place, and the arena was not cleaned between trials. This created a highly predictable environment, allowing scouts to retain information about previous trails and food locations. As a result, their movement was less random, and they consistently reached the intermediate distances where food was placed significantly faster than in the No-Food trials, where the arena was cleaned after each trial. The larger FPTs beyond 65 cm in the Food condition can be explained by a delay in long-distance exploration. This was due to the onset of the exploitation phase, during which scouts prioritized group recruitment rather than searching more distant regions of the arena.

The macroscopic differences observed in scouts' movement between the Food and No-Food conditions cannot be explained solely by local movement rules. Instead, they suggest that scouts are capable of incorporating past information (memory) into their exploratory behavior. This ability could account for the consistent discrepancies observed between the first food discovery times (TP1) in the experimental data and the simulations (see Fig. 5 and fig. S3). In the model, there is a strong stochastic effect across replicates, and no memory was included in the scouts' exploratory behavior. Altogether, these led to consistently longer TP1s compared to the experimental results, regardless of the proportion of scouts modeled.

### Proportion of scouts and recruits

We classified all trajectories occurring during the exploration phase into scouts and recruits and measured their relative proportions in each trial in the Food conditions. As shown in fig. S2, the results demonstrate a highly variable proportion of scouts and recruits across trials. In the model, we used a proportion of 0.5 for scouts, corresponding to the average proportion observed in the experiments.

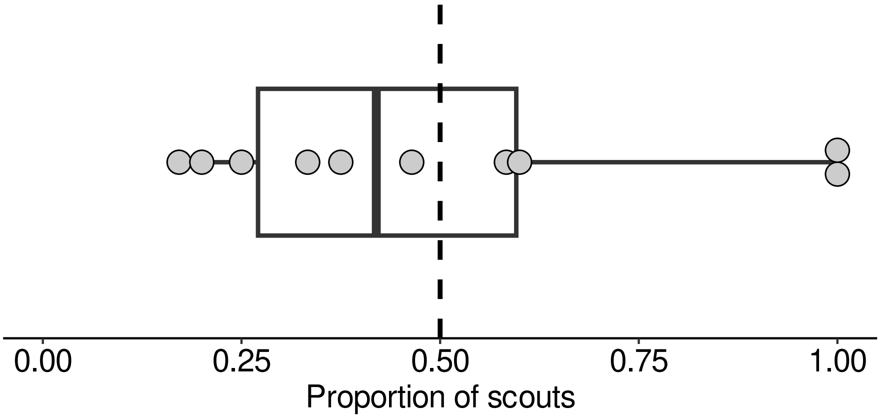


**Figure S2: Empirically estimated proportion of scouts**. We measured the relative proportion of scouts in each experimental trial (Food condition) during the exploration phase. We show the proportion for each replicate (dots), their distribution (box and whiskers), and the mean (dashed line) for all the replicates. In our model, we used the mean value (0.5) as the initial proportion of scouts.

### Foraging dynamics for different proportions of scouts

We carried out simulations with varying proportions of scouts to evaluate to which extent movement heterogeneity affects foraging dynamics. Increasing the proportion of scouts accelerated food discovery (TP1, dashed line in fig. S3). Despite this, TP1s were systematically overestimated in the simulations, likely because scouts learn from previous experiences (see supplementary text). However, the overall dynamics remained qualitatively similar, primarily driven by the exploitation phase and the food location information exchange process within the colony. Self-organized recruitment emerged from the interactions with a successful scout. Discovering food triggered an amplification cascade, driven by the increase in the value of information (encoded in $J_{ij}$, Eq. 1). This social feedback unfolded stochastically over time, producing a non-linear response in the colony, which was somewhat independent of the scout proportion.

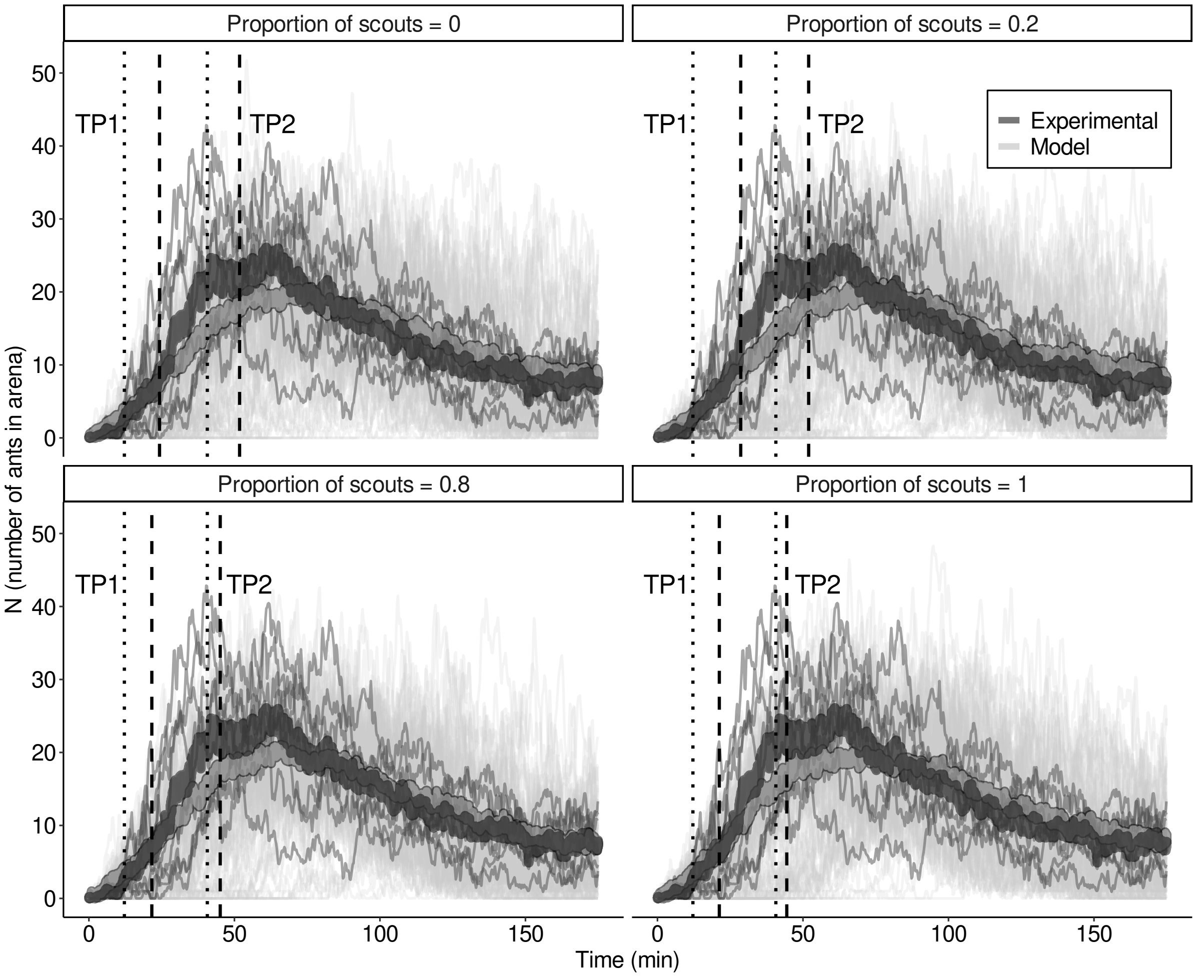


**Figure S3: Comparison of the foraging dynamics between 10 experimental trials (Food condition) and 100 stochastic simulations featuring different proportions of scouts**. The number of ants in the arena (y-axis) is shown for empirical (dark grey) and simulation (light grey) data. Thick lines represent the average, while thin lines in the background show each individual experiment or simulation. Dashed lines signify the first time food is detected (end of exploration phase, TP1) and the last time food is detected (end of exploitation phase, TP2) for the simulations. The dotted lines represent the same for the experimental trials. A, all recruits (0% of scouts). B, 80% recruits, 20% scouts. C, 20% recruits, 80% scouts. D, all scouts.

### First-passage times for scouts and recruits

To investigate the search potential of scouts and recruits, we simulated first-passage times (FPT) as a function of the distance from the nest for the two ant roles (fig. S4). Increasing the proportion of scouts decreased (FPT), suggesting that scouts are better at reaching far distances than recruits (fig. S4 inset). When comparing the limiting cases where all ants were either scouts or recruits (fig. S4), the differences in the FPT were statistically relevant even for small distances (18 cm and beyond). Notably, the variability is much higher for recruits, reflecting the broader range of trajectories they employ to cover a given spatial distance. In contrast, scouts exhibit consistently lower variability and are faster at reaching greater distances.

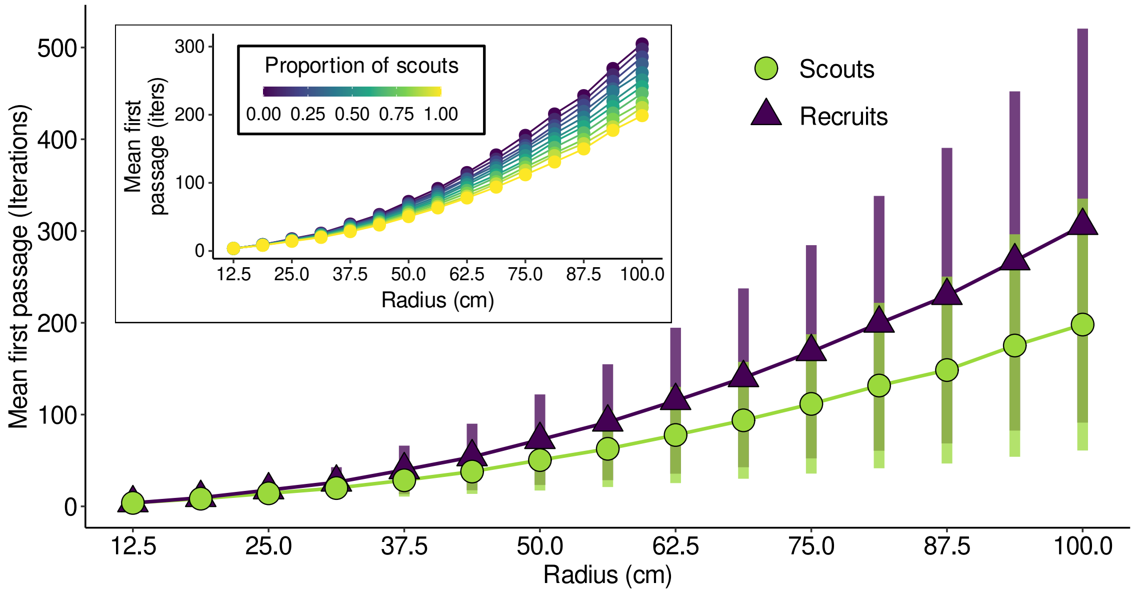


**Figure S4: First-passage times to assess macroscopic movement heterogeneity in scouts and recruits**. We show simulations of first-passage times (y-axis) in a population of 10000 ants as a function of distance-from-nest(x-axis). In the inset, we show how varying the proportion of scouts (being recruits the inverse of this proportion) affects the mean first-passage time of the population. The limit cases where all ants are either scouts (green) or recruits (purple) are displayed along with the standard deviation. Beyond R = 12.5 ($P = 0.076$), all observed differences are unlikely to happen by chance ($P \approx 0$, according to Wilcoxon tests).

### Scouts enable synchronized exploitation of multiple patches

We evaluated ants ability for parallel patch exploitation, examining how it varies with different proportions of scouts and levels of social copying. Specifically, we assessed the likelihood of multiple patches being exploited simultaneously and the duration of multi-patch exploitation (fig. S5). Our results indicate that social copying tends to work against patch parallelization. As social copying increases, both the probability (fig. S5A) and the duration overlap (fig. S5B) of simultaneous multi-patch exploitation decrease. Therefore, social copying drives ants to prioritize the exploitation of one patch before moving to another.

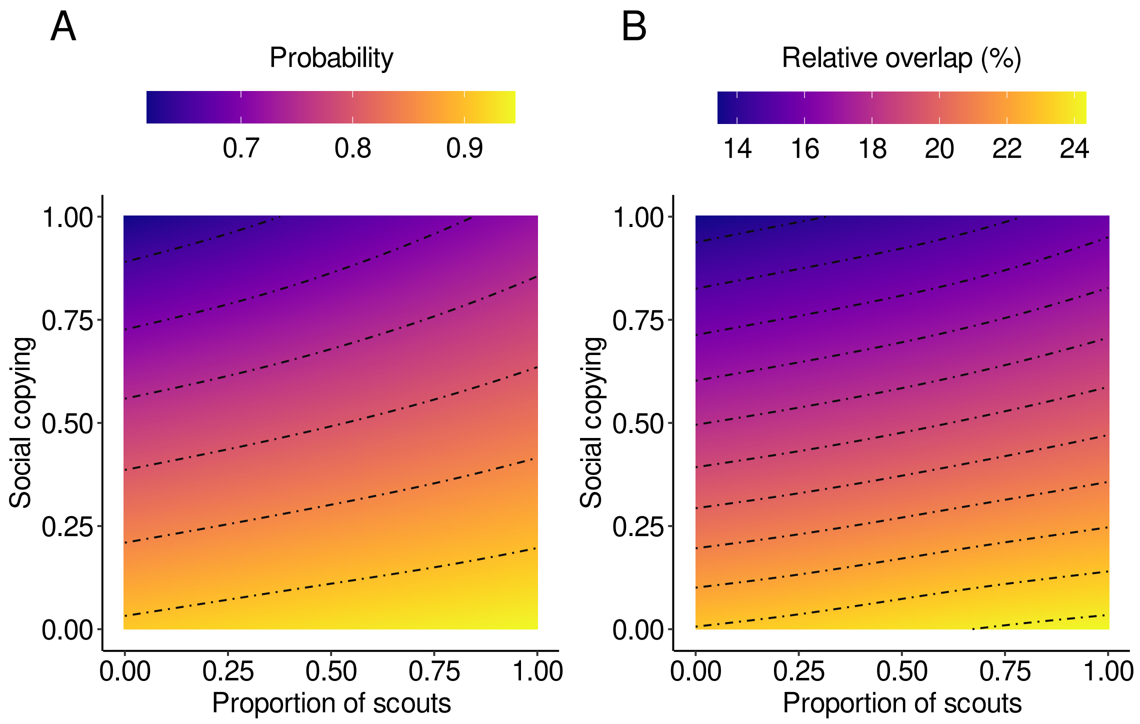


**Figure S5: Parallel exploitation of multiple food patches**. We assessed the synchronicity in the exploitation of multiple patches across varying proportions of scouts (x-axis) and social copying ants (y-axis). A, probability of parallel exploitation, measured as the proportion of simulations where the exploitation of both patches overlapped in time. B, duration of simultaneous exploitation relative to the total duration of the exploitation phase.

However, a higher proportion of scouts can help counter this effect. Increased scout numbers enhance the colony's search capacity, enabling scouts to locate unexploited patches while the rest of the colony focuses on exploiting a previously discovered patch. The scouts' propensity to disperse also keeps them from being drawn into the exploitation social feedback, allowing them to operate more independently. This behavior may benefit the colony by enabling the discovery of alternative patches, supporting overall foraging success.
